# Targeted therapy for *LIMD1*-deficient non-small cell lung cancer subtypes

**DOI:** 10.1101/2021.02.01.429178

**Authors:** Kathryn Davidson, Paul Grevitt, Maria F. Contreras G., Katherine S. Bridge, Miguel Hermida, Kunal M. Shah, Faraz K Mardakheh, Mark Stubbs, Paul A. Clarke, Rosemary Burke, Pedro Casado-Izquierdo, Pedro R. Cutillas, Sarah A. Martin, Tyson V. Sharp

## Abstract

An early event in lung oncogenesis is loss of the tumour suppressor gene *LIMD1 (LIM domains containing 1);* this encodes a scaffold protein, which suppresses tumourigenesis via a number of different mechanisms. Approximately 45% of non-small cell lung cancers (NSCLC) are deficient in LIMD1^1^, yet this subtype of NSCLC has been overlooked in preclinical and clinical investigations. Defining therapeutic targets in these LIMD1 loss-of-function patients is difficult due to a lack of ‘druggable’ targets, thus alternative approaches are required. To this end, we performed the first drug repurposing screen to identify compounds that confer synthetic lethality with LIMD1 loss in NSCLC cells. PF-477736 was shown to selectively target LIMD1 deficient cells *in vitro* through inhibition of multiple kinases, inducing cell death via apoptosis. Furthermore, PF-477736 was effective in treating LIMD1^−/−^ tumors in subcutaneous xenograft models, with no significant effect in LIMD1^+/+^ cells. We have identified a novel drug tool with significant preclinical characterization that serves as an excellent candidate to explore and define LIMD1-deficient cancers as a new therapeutic subgroup of critical unmet need.

**Significance Statement:** *Here we provide the first proof-of-concept data validating the scope for development of a targeted therapy against the non-small cell lung cancers (NSCLC) subtypes deficient in expression of the LIMD1 tumor suppressor gene. Approximately 45% of NSCLC are deficient in LIMD1^1^ representing at least 1.2 million lung cancer patients worldwide; yet this subtype has been ignored in preclinical and clinical investigations with no targeted therapies available. This seminal study applied synthetic lethality drug screening to target the loss/reduction of LIMD1 in lung cancer and normal cell lines, identifying and validating the multi-kinase inhibitor PF-477736 as a selectively cytotoxic compound towards LIMD1 deficient cells. This study provides rationale for further investigation into targeting LIMD1 loss in lung cancer, thereby addressing a critical unmet need for therapeutic approached to targeting LIMD1-deficent cancer subtypes.*

## Introduction

Lung cancer remains the most common cancer in the western world with ~2 million cases reported worldwide each year[1]. The most frequent type of lung cancer is non-small cell lung cancer (NSCLC), accounting for 84% of total cases, the majority of which are either lung adenocarcinomas (LUAD) or lung squamous cell carcinomas (LUSC)[1, 2]. The 5-year survival of lung cancer patients is only 19%, with minimal improvement in the past 30 years. Recent breakthroughs in immunotherapies and immune checkpoint PD-1/PD-L1 blockade for lung and several other cancers is encouraging [3–5]. Furthermore, advances in therapeutic research has led to the advent of highly specific targeted therapies such as tyrosine kinase inhibitors[6]. When used in combination with immunotherapy, this has achieved significant survival benefit for select patient subgroups[7]. However, only a proportion of patients will benefit (~10%-40% depending on cancer type), with the overall survival rates remaining largely unchanged[8–12]. This highlights the clear need for novel biomarkers and improved targeted therapies that extend beyond the current approaches, [13]. Furthermore, the plethora of mechanisms underlying lung cancer development and progression still remain largely unknown. Driver alterations have not yet been defined in ~40% of lung cancers. Although mutations in several well-known oncogenes and tumour suppressor genes have been detected in certain lung cancers with respect to their development and evolution, a large proportion of patients do not contain these common truncal mutations [14–17]. An improved understanding of the biology and enhanced treatment options are urgently needed.

*LIMD1* is a tumour suppressor gene encoded at the frequently ablated 3p.21.3 genomic locus in lung cancer. Reduced *LIMD1* copy number alterations in LUAD correlate with poor patient prognosis [18]. Furthermore, *LIMD1^−/−^* mice develop increased numbers and larger volumes of lung adenomas following exposure to the carcinogen urethane or upon crossing with *KRAS^G12D^* mice, highlighting loss of *LIMD1* as both a driver and major potential LUAD and LUSC susceptibility gene [18, 19]. Human lung cancers deficient in *LIMD1* expression represent 50% and 85% of LUAD and LUSC respectively [18], and have until now been almost completely overlooked. This biomarker signifies a new and exciting avenue for investigation in lung cancer biology and importantly with reference to this study, a novel treatment strategy for LIMD1-deficient tumours.

LIMD1 is a member of the Zyxin family of LIM-domain proteins, which feature three tandem LIM domains at the C-terminus that facilitate protein-protein interactions and an unstructured N-terminal pre-LIM region [20]. Whilst LIMD1 has no enzymatic function, it plays an important role in modulating many essential cellular homeostatic processes by operating as a nodal molecular scaffold [18, 21–29]. We have shown a critical role of LIMD1 as a core component of the microRNA-induced silencing complex (miRISC), scaffolding and dictating miRISC functional complexity [24], and in regulating the hypoxic response through mediating efficient degradation of HIF-1α by simultaneous binding of HIF prolyl-hydroxylases and the Von-Hippel Lindau protein (pVHL)[23, 30]. In addition, LIMD1 binds to and enhances the function of the retinoblastoma protein (pRB), thereby blocking E2F1-driven gene transcription and subsequent cell cycle progression[21]. Loss of LIMD1 and its multiple tumour suppressive functions, leads to alterations and disruption of these key homeostatic regulatory pathways, driving cellular transformation and cancer progression.

Despite *LIMD1’s* key homeostatic functions, level of ablation in LUAD/LUSC and the proportion of patients with this tumour subtype, there are currently no targeted therapies for LIMD1-deficient cancers. Defining therapeutic targets in these LIMD1 loss-of-function patients is difficult due to no clear ‘druggable’ enzymes identified that can be targeted, meaning alternative approaches are required. This is further complicated due to the number of diverse pathways impacted through loss of LIMD1, therefore targeting downstream pathways in isolation is not a feasible option. The concept of synthetic lethality provides a rationale for targeting tumour suppressor gene loss in cancer whereby cellular vulnerabilities acquired by cells following loss of tumour suppressors are exploited to induce cell death in tumour verses normal tissue. The prime example of this is the use of PARP inhibitors in *BRCA1* mutant cancers, which is now an approved targeted therapy in several cancers with loss of function *BRCA1* mutations [31].

To this end, we have performed the first proof-of-concept drug repurposing screen to identify synthetic lethal compounds with LIMD1 loss in lung cancer cells. Drug repurposing is an attractive option as a significant amount of preclinical data and safety profiling has already been generated for these compounds, allowing expedited clinical trials for alternative indications [32].

From our compound library screen, we identified a multi-kinase inhibitor, PF477736 that selectively kills LIMD1 negative cells compared to LIMD1 positive cells, whilst not affecting LIMD1 proficient cells. Whilst this inhibitor was designed as a checkpoint kinase 1 (Chk1) inhibitor, we have shown that Chk1 inhibition does not confer the synthetic lethal interaction with LIMD1 loss in these cells. Instead, our data indicates that this inhibitor affects a spectrum of kinases, inducing significant changes to the phosphoproteome specifically of LIMD1 negative cells. Finally, we show that this inhibitor of LIMD1 deficient cancer cell proliferation has therapeutic potential in lung adenocarcinoma, an aetiology of importance in LIMD1 biology. This study provides the first proof-of-concept that LIMD1 expression can be used as a stratification marker for treatment, identifying a large group of lung cancer patients that could benefit from a targeted therapy against *LIMD1* loss.

## Results

### Drug repurposing screen has identified PF477736 to selectively inhibit LIMD1^−/−^ cells

To identify compounds that selectively target LIMD1^−/−^ cells, we screened CRISPR-Cas9 generated isogenic LIMD1^+/+^ and LIMD1^−/−^ HeLa cells with a compound library of 485 molecularly targeted small molecules. This drug library was collated to include FDA-approved drugs, clinical candidates and compounds against known cancer pathways [33, 34]. Cells were treated with a 1 μM concentration of each compound, and cell viability was determined after five days. Upon determination of the ΔZ-scores we identified the Checkpoint Kinase 1 (Chk1) inhibitor PF-477736 as our lead hit, such that it had one of the highest ΔZ-scores, causing significantly decreased cell viability in LIMD1^−/−^ cells, compared to the LIMD1^+/+^ cells, as well as not showing overt toxicity in the LIMD1^+/+^ line ***(Fig. 1A).*** We validated this synthetic lethal interaction across a range of concentrations, in multiple LIMD1^−/−^ cell clones ***(Fig. 1B-C).*** Of note, this phenotype was validated in our isogenic pair of CRISPR-Cas9 generated lung adenocarcinoma A549 LIMD1^+/+^ and LIMD1^−/−^ cells ***(Fig. 1B-C)*** indicating this effect was not cell line specific and was relevant in the context of lung cancer biology. In both LIMD1 isogenic cell models, there was a ~2-fold selectivity towards LIMD1^−/−^ cells compared to LIMD1^+/+^ controls ***(Fig. 1D-E)*** demonstrated by a significant difference in SF50 values ***(Fig. S1A-B).*** We further validated this effect using long term clonogenic drug assays, which showed a significant decrease in the number of colonies for LIMD1^−/−^ cells compared to the LIMD1^+/+^ upon PF-477736 treatment ***(Fig. 1F-G).*** In addition, we treated these isogenic lines with 1 *μ*M PF4 and measured cell proliferation using bright field imaging on Incucyte Zoom; whilst there was a modest reduction in proliferation in the LIMD1^+/+^ lines upon PF4 treatment (~1.6-fold reduction in AUC), this effect was significantly more selective in both LIMD1^−/−^ clones (~7-fold reduction in AUC) *(**Fig. 1H)**.* Notably, PF-477736 treatment induced ‘blebbing-like’ structures of cell membranes in LIMD1^−/−^ cells, suggesting that cells may be undergoing increased apoptosis ***(Fig. S1C).*** This was confirmed by increased PARP cleavage in LIMD1^−/−^ clones compared to LIMD1^+/+^ controls upon PF-477736 treatment. ***(Fig. 1I-J)***. Annexin V staining identified increased early and late apoptotic cell populations in PF-477736 treated LIMD1^−/−^ cells ***(Fig. S1D)***. Taken together, these data indicate that PF-477736 treatment can selectively target LIMD1^−/−^ cells, identifying apoptosis as the dominant mechanism of cell death.

**Figure 1.**
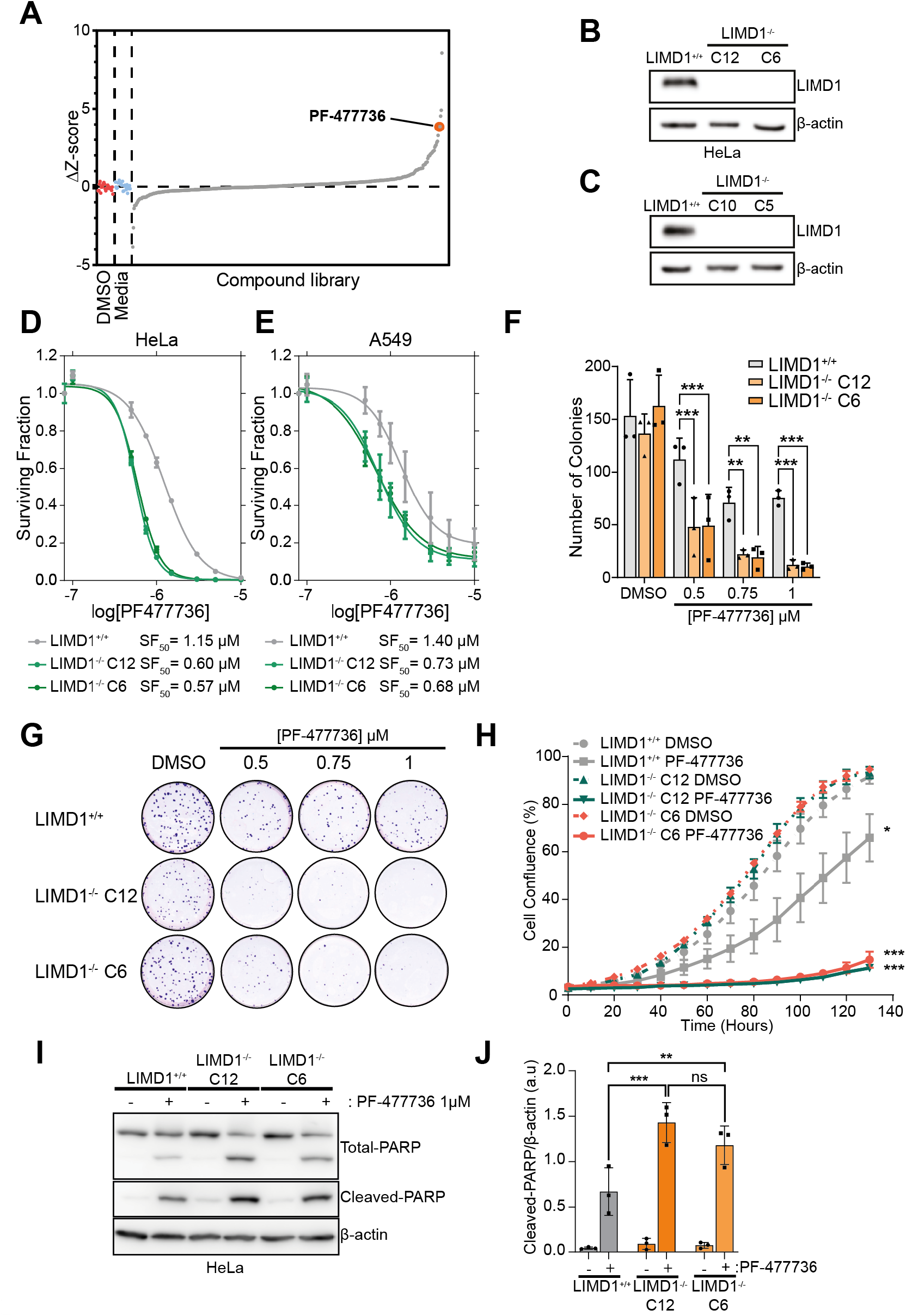
PF-477736 is a selective inhibitor of LIMD1 deficient cells. **A)** Waterfall plot of DZ-Scores from compound library screen. Isogenic LIMD1^−/−^ and control lines were treated with a 1 mM dose and cell viability measured after 4 days (n = 3). **B-C)** Immunoblot of CRISPR-Cas9 generated LIMD1^−/−^ cell lines in HeLa and A549. **D-E)** Dose response curves of PF-477736 in A549 and HeLa isogenic LIMD1^−/−^ lines. Cells were drugged twice over 4 days before measuring cell viability and calculating Surviving Fraction (n = 3). **F)** Growth of HeLa isogenic LIMD1^−/−^ measured using Incucyte Zoom following 1 mM treatment with PF477736 (n = 3; one-way ANOVA comparing AUC for each curve). **G-H)** Colony formation assay of HeLa isogenic LIMD1^-^ cells following treatment of PF-477736 for 10 days. Cells were treated every 2-3 days with indicated concentration of PF-477736 before fixation and staining (n = 3, two-way ANOVA). **I-J)** Western blot densitometry quantifying PARP cleavage in HeLa isogenic LIMD1^−/−^ lines treated with PF-477736 for 48 hours. (n = 3, two-way ANOVA). ns p>0.05, *p≤0.05, **p≤0.01, ***p≤0.001.

### PF-477736 selectively kills LIMD1^−/−^ cells independent of Chk1 inhibition

PF-477736 was originally developed by Pfizer for use in combination therapy with DNA damage inducing agents as a sub-nanomolar ATP-competitive inhibitor of checkpoint kinase 1 (Chk1) (Chk1 Ki = 0.49 nM)[35]. However, neither of the two other Chk1 inhibitors in the compound library (AZD7762 and LY2603618) were identified as hits in the compound screen ***(Fig S2A).*** We reasoned this may have been an artefact of the single concentration dose used in drug screening. We tested an alternative Chk1 inhibitor, SCH900776 (MK-8776, Chk1 Ki = 3 nM), to establish if this caused LIMD1^−/−^ specific cell death ***(Fig. 2A-B).*** We did not observe any difference in cell viability between our HeLa isogenic lines following treatment. RNAi mediated transient knockdown of *CHEK1* in our isogenic lines was also performed, and once again did not induce any differences in cell viability between these lines ***(Fig. 2C-D).*** These data indicate that Chk1 inhibition is not synthetically lethal with LIMD1 loss, and therefore the effects we see with PF477736 are likely to be through inhibition of an alternative kinase or kinases.

**Figure 2.**
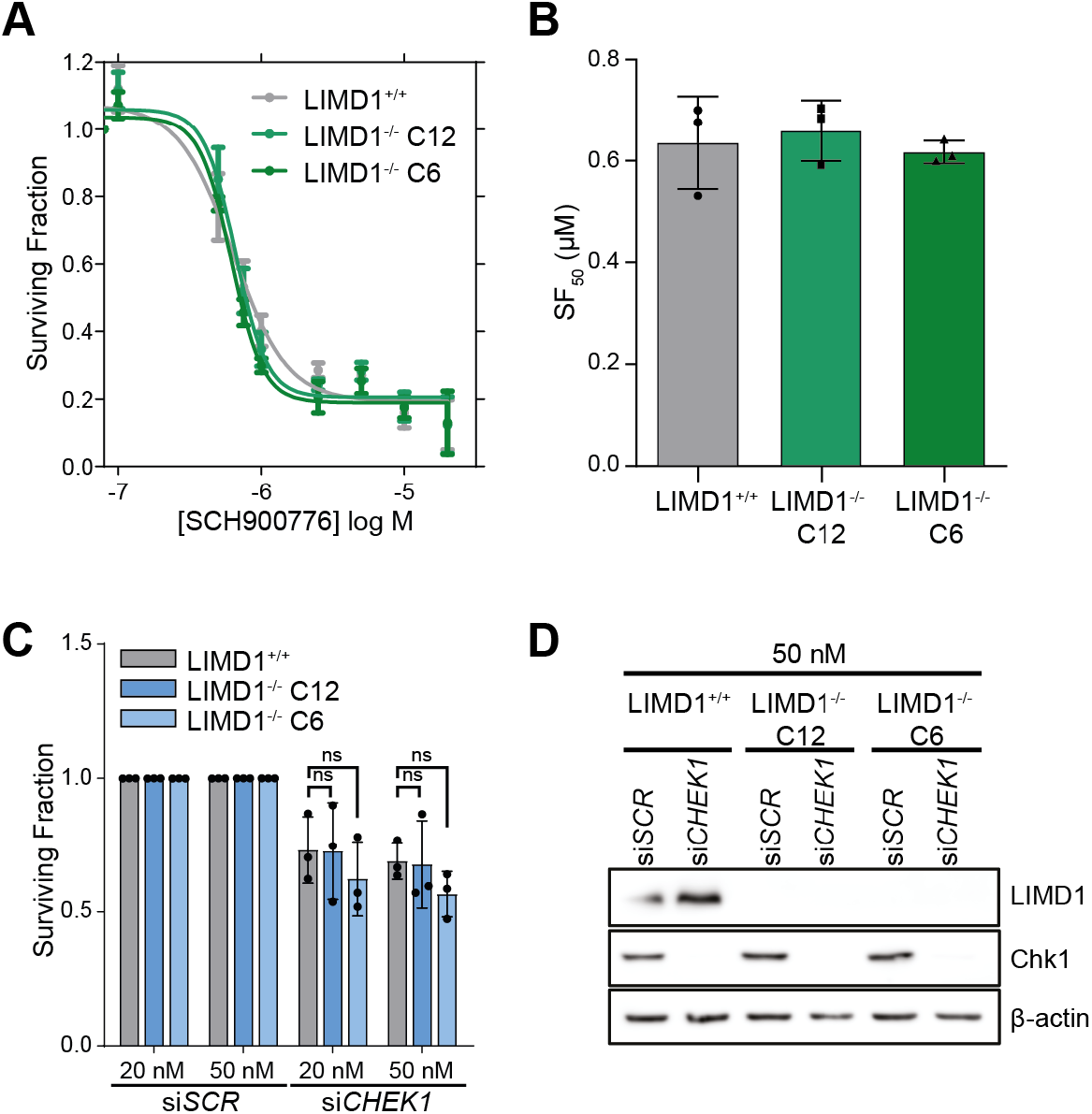
PF-477736 selectively kills LIMD1^−/−^ cells independent of Chk1 inhibition. **A)** Dose response curve of SCH900776 in HeLa isogenic LIMD1^−/−^ lines. Cells were treated for 4 days before measuring viability and calculating surviving fraction (n = 3). **B)** Bar chart of SF50 values from panel A (n = 3, one-way ANOVA). **C)** Surviving fraction of HeLa isogenic LIMD1^−/−^ lines transfected with siRNA against *CHEK1* at 50 nM and 20 nM for 72 hours (n=3, two-way ANOVA) ns p>0.05. **D)** Immunoblot of Chk1 and LIMD1 in HeLa isogenic LIMD1^−/−^ lines transfected with siRNA against *CHEK1* (20nM, 72 hours) (n=3).

### PF-477736 is a broad-spectrum kinase inhibitor that elicits LIMD1^−/−^ specific cellular changes in the phosphoproteome

With the aim of identifying the kinase responsible for synthetic lethality with LIMD1 loss upon PF-477736 treatment, we tested the activity of PF-477736 on a panel of 403 recombinant protein non-mutant kinases and 59 clinically relevant disease mutant kinases. For this, we utilised the DiscoverX KINOMEscan platform, which involves an *in vitro* ATP-independent competition assay to measure kinase activity [36]. Optimum inhibitor concentration is defined as 3-10 fold higher than the Ki of targeted interactions, therefore as we observe the strongest synthetic lethal interaction at 1 μM, we opted to perform profiling at a PF-477736 concentration of 3 μM. Surprisingly, 303 out of the 468 kinases tested were inhibited by over 50%, upon PF-477736 treatment compared to the DMSO control ***(Fig. 3A).*** We observed a wide range of inhibition across the kinases tested, with the activity of 18.4% of kinases inhibited by 99%, further highlighting the broad specificity of this inhibitor ***(Fig. 3B).*** A percentage of control value of <1% indicates a Kd value of <30nM., Chk1 activity was reduced to 0.45% by PF-477736, and a number of other kinases are more potently inhibited by this drug ***(Fig. 3C).*** Next, in an endeavour to elucidate which kinases are inhibited by PF-477736 in the cell as opposed to the *in vitro* study, we analysed the phosphoproteome in our isogenic lines upon drug treatment ***(Fig. 3D).*** Following 1-hour treatment with PF477736 we did not observe any significant changes in the phosphoproteome of HeLa LIMD1^+/+^ cells; strikingly however, there were numerous significant changes occurring in the HeLa LIMD1^−/−^ cells ***(Fig. 3D).*** Principle component analysis (PCA) on this data showed no clear separation between treated and untreated samples in the LIMD1^+/+^ cell line, but a clear separation in the LIMD1^−/−^ cell line **(Fig. S3A-B)**. This result emphasizes that PF-477736 treatment elicits cellular phosphorylation changes specifically upon loss of LIMD1 expression. We next utilised Kinase-Substrate Enrichment Analysis (KSEA) to infer kinase activities from our quantitative phosphoproteomics data [37]. This analysis identified a number of kinases (CK2A1, CDK1, PKCA, ERK1 and Akt1) that were significantly more active in LIMD1^−/−^ cells compared to LIMD1^+/+^ controls ***(Fig. S3C).*** Furthermore, CK2A1, PKCA and Akt1 were all significantly inhibited following PF4 treatment specifically in LIMD1^+/+^ cells. Interestingly, LIMD1^+/+^ cells exhibited increased CK2A1 activity after treatment ***(Fig. 3E).*** In our DiscoverX KINOMEscan data these kinases were inhibited by 90.1%, CK2A1; 89%, PKCA and 77%, Akt1 ***(Fig. 3A-C).*** To determine whether loss of these kinases alone could induce synthetic lethality upon LIMD1 loss, we knocked down expression (via siRNA) of each kinase individually or in combination, however, there were no significant changes in cell viability between our isogenic lines with each of these targeted knock-downs ***(Fig. S3D).*** We next reasoned that knocking down kinase levels significantly with siRNA could possibly still leave very low levels of protein and activity that could still maintain viability and thus opted to examine small molecule inhibitors against the targets (both individually and in combination at a dose that corresponds with SF_80_) with high efficacy of inhibition for the indicated kinases/pathways. This drug targeted approach was based on inhibitors: MK-2206, IC_50_s of 5 nM, 12 nM, and 65 nM for AKT1, AKT2, and AKT3; SCH900776 (MK-8776); Silmiterasertib with an IC_50_ of 1 nM for Casein kinase 2 and Gouml 6983 (Go-6983), a potent broad spectrum PKC inhibitor with IC_50_ values are 7, 7, 6, 10, 60 and 20000 nM for PKCα, PKCβ, PKCγ, PKCδ, PKCζ and PKCμ respectively. These drug analyses and combinations did not show any selectivity towards LIMD1 deficient cells ***(Fig. S3E-F).*** This suggests that other pathways, targets or even non-enzymatic susceptibilities are involved, or indeed far greater integrated complex targeting susceptibilities. Regardless of not identifying a specific kinase and/or pathway, these data do indicate that loss of LIMD1 increases activity and dependency on a panel of kinases including CK2A1, PKCA and Akt1 which were inhibited by PF4 treatment, leading to increased apoptosis in LIMD1^−/−^ cells.

**Figure 3.**
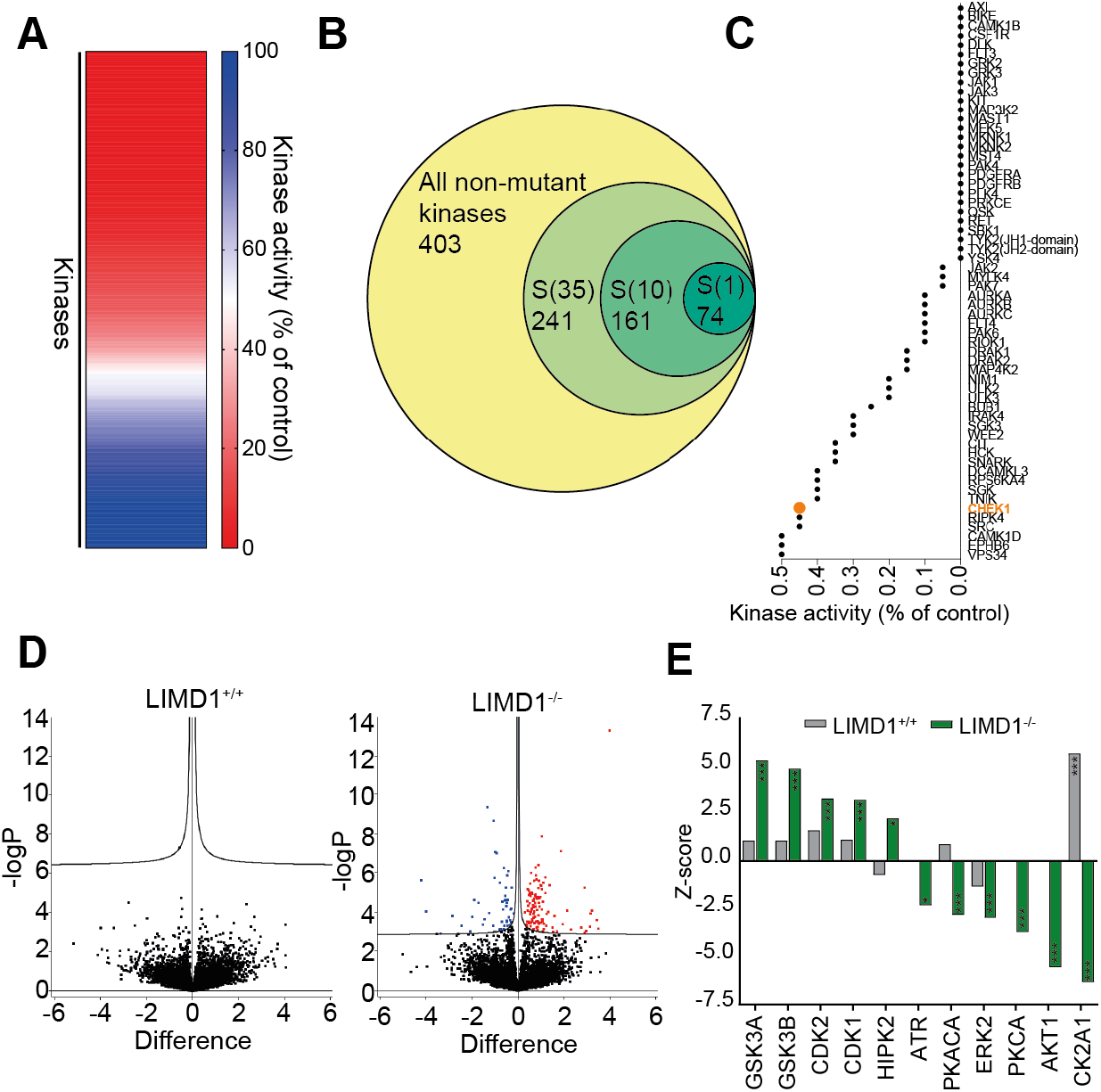
PF-477736 is a broad-spectrum kinase inhibitor that elicits LIMD1^−/−^ specific cellular changes in the phosphoproteome. **A)** Heatmap showing remaining kinase activity of a panel of in vitro kinases upon 3 mM PF477736 treatment in DiscoverX KINOMEscan assay. **B)** Venn diagram representing the proportion and number of kinases inhibited to less than 35%, 10% and 1% of control activity. **C)** Waterfall plot of kinases most inhibited by PF477736 (3 mM) in the *in vitro* kinase assay. **D)** Volcano plot of phosphosite changes between 1μM PF477736 treated and DMSO control lysates in isogenic HeLa lines. Cells were harvested following 1 hour drug treatment. No significant phosphosite changes were induced by PF477736 in the LIMD1^+/+^ cell line, compared with 54 reduced and 119 increased phosphosites in the LIMD1^−/−^ cell line. Cut off point for statistically significant phosphosite is a false discovery rate of > 0.05 and s0 of 0.01. (n = 3). **E)** Kinase Substrate Enrichment Analysis (KSEA) of phosphoproteomics shows kinases significantly affected by PF477736 treatment in LIMD1^+/+^ (grey) or LIMD1^−/−^ cells. *p≤0.05, **p≤0.01, ***p≤0.001

### PF-477736 treatment represents the first proof-of-concept for targeted inhibition of LIMD1 deficient lung cancers

Previous work from our group has extensively characterised the role of LIMD1 as a tumour suppressor in lung cancer, therefore we investigated the potential of PF-477736 as a therapeutic against LIMD1 deficient lung cancer cell lines. We have already shown selectivity of PF-477736 against LIMD1^−/−^ A549 cells, a NSCLC adenocarcinoma cell line ***(Fig. 1E).*** To test PF-477736 in non-transformed lung cells we generated LIMD1 hypomorphic mutant Small Airway Epithelial Cells (SAEC) that had been immortalised by overexpressing Bmi1 (SAEC-Bmi1). These SAEC are a mixture of both type I and type II alveolar cells, thereby serving as an appropriate model of lung adenocarcinoma progenitor cells, which have been CRISPR-Cas9 edited and express a N-terminal truncated form of LIMD1 in significantly lower levels compared to non-targeting controls, thereby representing an *in vitro* model of early LIMD1 ablation/loss in the development of adenocarcinoma ***(Fig. 4A).*** Comparable with our other isogenic cell lines, we observed ~2-4-fold selectivity for SAEC LIMD1^−/−^ cells ***(Fig. 4B, S4A).*** Next, we treated a panel of lung adenocarcinoma cell lines exhibiting a range of LIMD1 protein expression, with PF-477736 ***(Fig. 4C-D).*** We observed a significant positive correlation (Pearson’s correlation coefficient = 0.579, p= 0.0302) with LIMD1 protein expression and surviving fraction at 1μM thereby indicating that it may be possible to use LIMD1 expression as a biomarker to stratify patients for targeted therapy treatment efficacy ***(Fig. 4D).*** To test the efficacy of PF-477736 *in vivo* we inoculated NOD/SCID mice with our A549 isogenic lines subcutaneously and treated with PF477736 twice on indicated days. Unexpectedly, we observed decreased tumour growth in the LIMD1^−/−^ tumours, which may represent difference between cell fitness during initial tumour implantation, as relative tumour growth was similar across lines (***Fig S4A-B).*** Importantly, LIMD1 expressing tumours (LIMD1^+/+^) were unaffected by PF-477736 treatment *in vivo,* however we observed a significant decrease in tumour growth in the LIMD1^−/−^ tumours upon treatment ***(Fig. 4E, S4B).*** Staining of these tumours with markers for cell proliferation (Ki67, ***Fig. 4F)*** and apoptosis (cleaved Caspase-3, ***Fig. 4G),*** revealed that PF-477736 selectivity inhibits proliferation in LIMD1-deficient lung xenografts and increases apoptosis within these tumours, in agreement with our *in vitro* data.

**Figure 4.**
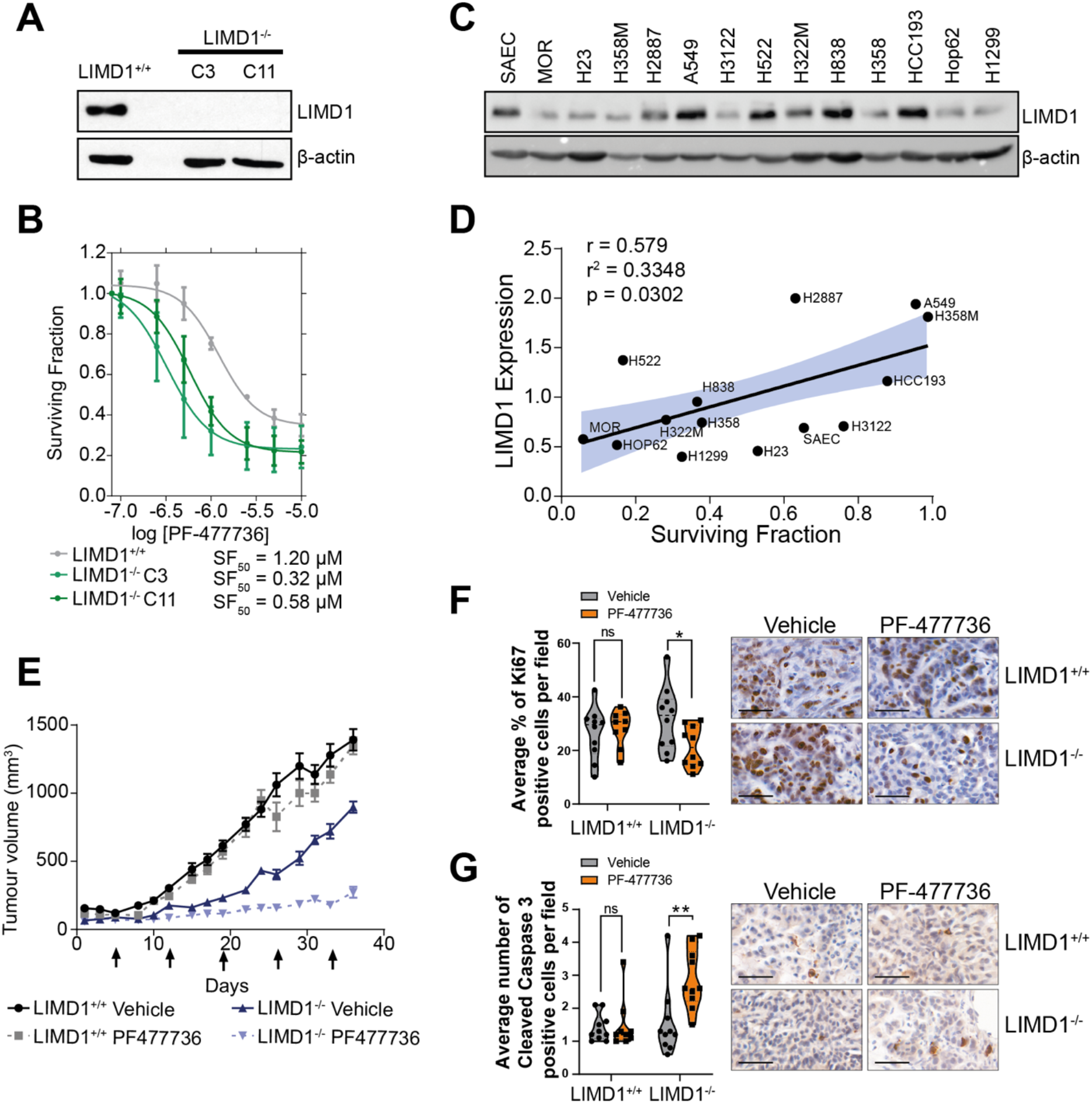
PF-477736 treatment is proof-of-concept inhibitor of LIMD1 deficient lung cancers. **A)** Immunoblot of isogenic LIMD1*mut* Small Airway Epithelial cells and control. **B)** Dose response of PF-477736 in isogenic LIMD1*mut* Small Airway Epithelial cells. Cells were treated twice prior to measuring cell viability and calculating surviving fraction (n = 3). **C)** Immunoblot of LIMD1 in a panel of lung adenocarcinoma cell lines (representative blot from n = 3). **D)** Pearson’s correlation coefficient between LIMD1 protein expression and surviving fraction of indicated cell line after treatment with 1 mM PF477736. **E)** Tumour growth of subcutaneous A549 isogenic LIMD1^−/−^ xenografts implanted into the flank of NOD/SCID mice. Mice were treated twice on indicated days with vehicle or PF477736 (7.5 mg/kg) (n = 10 per group). **F-G)** Immunohistochemical staining and scoring of Ki67 (**F**) and Cleaved Caspase-3 (**G**) in mouse xenograft tumours. (n = 10 per group, two-way ANOVA)

To summarise our results, we have identified that the multi-kinase inhibitor PF-477736, by targeting multiple susceptibility pathways, selectively induces apoptosis in LIMD1^−/−^ cells, whilst sparing LIMD1^+/+^ cells. This is the first evidence supporting a targeted therapeutic approach for the treatment of a large proportion of lung cancers with LIMD1 loss.

## Discussion

Lung cancer has a staggering disease burden, with a clear need for novel targeted therapies for larger patient populations. Loss of the tumour suppressor gene *LIMD1* is an attractive therapeutic option as we have identified copy number alterations in ~47% of lung adenocarcinoma patients [18]. Here we show a proof-of-concept study identifying PF-477736 as a selective inhibitor of LIMD1 deficient cells ***(Fig1).*** Crucially, we observe a correlation with LIMD1 expression and drug sensitivity, indicating that we may be able to use LIMD1 expression as a biomarker for new targeted therapies and thus indicate potential responsive treatment efficacies.

PF477736 is a checkpoint kinase 1 (Chk1) inhibitor, however our results indicate a number of alternative kinases are potently inhibited *in vitro* and *in cellulo* by PF-477736 ***(Fig. 3).*** This finding is perhaps not surprising, as an expanding number of studies have shown that several ATP-pocket kinase inhibitors which have been used in clinical trials, exhibit off-target mechanisms of action distinct from the primary drug target [38]. Our data indicates potential targeting of multiple pathways is required for the treatment of LIMD1 deficient disease; previous work from our group and others have highlighted a number of pathways that LIMD1 plays a crucial role in regulating [24, 39]. Therefore, it is unsurprising that targeting just one LIMD1-regulated pathway is insufficient to recapitulate PF-477736 induced cell death. Broad spectrum inhibitors have been crucial for targeting cancers, such as dual inhibition of PDGFRa and FGFR2 by pazopanib in SMARCB1 deficient rhabdoid tumors [40, 41]. This fits within our model of LIMD1 being a nodal gene, loss of which re-orchestrates numerous cellular pathways which requires inhibition of multiple kinases in order to induce cell death. For our analysis we focused on protein kinases as PF-477736 was designed as an inhibitor of the ATP moiety to block kinase function, however, there remains the possibility that PF-477736 is acting on a non-kinase enzyme or a kinase not covered by our analysis.

This proof-of-concept study has identified, for the first time, a selective inhibitor of LIMD1 deficient cells. We have shown that it is possible to target these cells with a small molecule, allowing for the potential targeted treatment of a large proportion of lung cancer patients with LIMD1 deficient tumours. Targeting such deficient cancers is imperative, as LIMD1 loss has been further observed in breast, cervical, gastric, renal and head and neck cancers [42–47]. Our study has identified a novel therapeutic strategy based on LIMD1 status which exploits the loss of this tumour suppressor, offering the potential for targeted treatment for this large cancer patient population and significantly reduce disease burden worldwide.

## Materials and Methods

### Cell Culture

Cells were maintained in DMEM (HeLa, A549) or RPMI (HCC193, H1299, Hop62, H358, H838, H358M, H2887, H522, H322M, H23, MOR, H3122) (Sigma) supplemented with 10% FCS and 1% Penicillin/Streptomycin solution in a humidified 37°C incubator with 5% CO_2_. SAEC were maintained in complete Airway Epithelial Cell Medium (ATCC). SAEC cells were immortalised using pFLRu-Bmi-1. Al cell cultures are regularly tested for mycoplasma.

### Drug screening

Cells were plated in 96-well plates at an optimised cell density, and after 24 hours treated with vehicle (0.01% DMSO) or the compound library at a final concentration of 1uM. Cells were dosed again after 48 hours. Cell viability was assessed using CellTitre-Glo (Promega) after 4 days of drug exposure, according to manufacturer’s instructions. Luminescence readings from each well were log transformed and normalized according to the median signal on each plate and then standardized by Z-score statistic, using the median absolute deviation to estimate the variation in each screen. Z-scores were compared to identify compounds that cause selective loss of viability in LIMD1^−/−^ cells compared to LIMD1^+/+^ cells.

### Drug validation experiments

PF477736 was purchased from Sigma-Aldrich. SCH900776, MK-2206 Go-6983, Silmitasertib was purchased from Selleckchem. In line with drug screen protocol, cell were seeded into a 96-well plate as treated 24 hours later with a concentration range of drug, or DMSO control. Cells were re-dosed with drug after 48 hours and cell viability determined after 4 days of drug exposure using CellTitre Glo. MK-2206, Go-6983 and Silmitasertib were purchased from Sigma-Aldrich. For combination studies, SF_80_ was calculated using the formula SF_80_ = (80/(100-80)^√Hill slope^ × SF_50_. Cells were plated and dosed as above before measuring viability with CellTitre Glo.

### Colony formation assay

Cells were seeded at a low, colony forming density in a 6-well plate. 24 hours post seeding, cells were treated with drug or vehicle control. Drug containing media was refreshed every 2-3 days and cells were fixed in methanol after 10 days. Colonies were stained with 0.05% crystal violet and counted by eye.

### Incucyte

Growth curves were generated using the IncucyteZOOM live-cell imaging platform (Essen Bioscience). Cells were seeded into 96-well plates and were drugged 24 hours later with either 1 μM PF477736 or DMSO control. Images were captured at 10x magnification on the Incucyte every 2 hours. Drug was refreshed after 68 hours. Cell confluence per well was calculated using the IncucyteZOOM software (Essen Bioscience).

### Protein analysis

Cell pellets were lysed in RIPA buffer (150nM NaCl, 1% (v/v) IGEPAL, 0.5% (w/v) Deoxycholic acid, 0.1% (w/v) SDS, 50mM Tris) supplemented with Protease and Phosphatase inhibitors. Protein was quantified using the Pierce BCA protein assay kit (Thermo Fisher Scientific). Lysates were electrophoresed on acrylamide gels of appropriate acrylamide percentage, transferred onto PVDF membranes and immunoblotted using the following antibodies: LIMD1 (in house), B-actin (Sigma), Total PARP (CST), Cleaved PARP (CST), Chk1 (CST). Anti-IgG horseradish peroxidase (Dako) and chemiluminescent detection (Thermo Fisher Scientific) were used to develop immunoblots.

### siRNA transfections

siRNA targeting Chk1, CSNK2A1, AKT1, PKCA, PLK1 and non-targeting control were obtained as SMARTPools from Dharmacon. siRNAs were transfected using Lipofectamine RNAiMax (Invitrogen) according to manufacturer’s instructions. 96-well plates were used for cell viability endpoints, and 12-well or 6-well plates were used for protein extraction for determination of protein knockdown by immunoblot.

### Annexin V staining

Cells were seeded into 10cm dishes at a density of 1×10^4^ cells/dish. 24 hours later dishes were treated with 1uM PF477736 or DMSO vehicle control. After 48 hours, cells were trypsinised and harvested (including those in media and PBS washes), counted and 1×10^6^ cells resuspended into 1 annexin-binding buffer (Thermo Fisher Scientific). 2ul Alexa Flour 488 Annexin V (Thermo Fisher Scientific) and 1ul 100ug/ml PI were added to each 100ul of cell suspension, and incubated for 15 minutes. Single stained and unstained controls were stained accordingly. Samples were run through the BD LSR Fortessa flow cytometer (Becton Dickinson, USA), recording 10,000 events for each sample. Data was analysed using FlowJo version 10 (FlowJo LLC).

### Kinase profiling

Kinase profiling was conducted by DiscoverX (USA). PF-477736 was dissolved in DMSO and diluted to 3mM (1000x screening concentration) for shipment. Profiling was performed by Discover X as per their in-house protocols.

### Phosphoproteomics

Mass spectrometry analysis was conducted as described by Casado *et al.* (2018)[48]. Briefly, HeLa LIMD1^+/+^ and LIMD1^−/−^ were seeded into 10 cm dishes at 7 x 10^5^ cells/dish. 48 hours post-seeding dishes were treated with either 1 μM PF477736 or DMSO vehicle control for 1 hour. Cells were washed 3x in ice-cold PBS containing protease and phosphatase inhibitors (1 mM NaF and 1 mM Na_3_VO_4_). Dishes were lysed in 200 μl of lysis buffer (8M Urea in 20 mM HEPES (pH 8.0) supplemented with, 1 mM Na_3_VO_4_, 1 mM NaF, 1 mM β-glycerol phosphate and 2.5 mM Na_2_H_2_P_2_O_7_).) and cells scraped and transferred into Protein Lo-bind tubes (Eppendorf). Samples were sonicated for 10 cycles (30s on & 40s off) in a Diagenode Bioruptor® Plus. Samples were centrifuged at 20,000 xg for 10 minutes, 4°C and supernatant transferred to 1.5 ml Protein Lo-bind tubes. BCA assay was conducted to quantify protein concentration.

Protein suspensions of 400 μg of protein in a volume of 200 μL were subjected to cysteine reduction and alkylation using sequential incubation with 10 mM dithiothreitol (DDT) and 16.6 mM iodoacetamide (IAM) for 1 h and 30 min, respectively, at 25°C with agitation. The urea concentration in the protein suspensions was reduced to 2 M by the addition of 600 μL of 20 mM HEPES (pH 8.0) and 100 μL of equilibrated trypsin beads were added and samples were incubated overnight at 37°C. Trypsin beads (50% slurry of TLCK-trypsin) were equilibrated with 3 washes with 20 mM HEPES (pH 8.0). The following day, trypsin beads were removed by centrifugation (2,000 xg at 5°C for 5 min) and samples were desalted using Oasis HLB cartridges (Waters).

Briefly, cartridges set in a vacuum manifold device, with a pressure adjusted to 5 mmHg, were conditioned with 1 mL acetonitrile (ACN) and equilibrated with 1.5 mL of wash solution (0.1% trifluoroacetic acid (TFA), 2% ACN). Then, peptide solutions were loaded into the cartridges and washed twice with 1 mL of wash solution. Peptides were eluted with 0.5 mL of glycolic acid buffer A (1 M glycolic acid, 5% TFA, 50% ACN).

For phosphopeptide enrichment, eluents were normalized to 1 mL with glycolic acid buffer B (1 M glycolic acid, 5% TFA, 80% ACN) and incubated with 25 μl of TiO_2_ solution (500 mg TiO_2_ beads in 500 μL of 1% TFA) for 5 min at room temperature. TiO_2_ beads were packed by centrifugation into empty spin columns previously washed with ACN. TiO_2_ bead pellets were sequentially washed by centrifugation (1500 xg for 3 min) with 100 μL of glycolic acid buffer B, ammonium acetate buffer (100 mM ammonium acetate in 25% ACN) and twice with neutral solution (10% ACN). Spin tips were transferred to fresh tubes and phosphopeptides were eluted by adding 50 μL of elution solution (5% NH_4_OH, 7.5% ACN) and centrifuging the spin-tips at 1500 xg for 3 min. This elution step was repeated a total of 4 times. Finally, samples were frozen in dry ice for 15 min, dried in a SpeedVac vacuum concentrator and stored at −80°C.

Peptide pellets were reconstituted in 13 μL of reconstitution buffer (20 fmol/μL enolase in 3% ACN, 0.1% TFA) and 5 μL were loaded twice onto an LC-MS/MS system consisted of a Dionex UltiMate 3000 RSLC coupled to Q Exactive™ Plus Orbitrap Mass Spectrometer (Thermo Fisher Scientific) through an EASY-Spray source. Chromatographic separation of the peptides was performed using the mobile phases A (3% ACN; 0.1% FA) and B (99.9% ACN; 0.1% FA). Peptides were loaded in a μ-pre-column and separated in an analytical column using a gradient running from 3% to 23% B over 60 min. The UPLC system delivered a flow of 2 μL/min (loading) and 250 nL/min (gradient elution). The Q Exactive Plus operated a duty cycle of 2.1s. Thus, it acquired full scan survey spectra (m/z 375–1500) with a 70,000 FWHM resolution followed by data-dependent acquisition in which the 15 most intense ions were selected for HCD (higher energy collisional dissociation) and MS/MS scanning (200–2000 m/z) with a resolution of 17,500 FWHM. A dynamic exclusion period of 30s was enabled with m/z window of ±10 ppm.

Peptide identification was automated using Mascot Daemon 2.6.0. Thus, Mascot Distiller v2.6.1.0 generated peak list files (MGFs) from RAW data and Mascot search engine (v2.6) matched the MS/MS data stored in the MGF files to peptides using the SwissProt Database (SwissProt_2016Oct.fasta). Searches had a FDR of ~1% and allowed 2 trypsin missed cleavages, mass tolerance of ±10 ppm for the MS scans and ±25 mmu for the MS/MS scans, carbamidomethyl Cys as a fixed modification and PyroGlu on N-terminal Gln, oxidation of Met and phosphorylation on Ser, Thr, and Tyr as variable modifications.

A label free procedure based on extracted ion chromatograms (XICs) quantified all Identified peptides. Missing data points were minimized by constructing XICs across all LC-MS/MS runs for all the peptides identified in at least one of the LC-MS/MS runs [49]. XIC mass and retention time windows were ±7 ppm and ±2 min, respectively. Quantification of peptides was achieved by measuring the area under the peak of the XICs. Individual peptide intensity values in each sample were normalized to the sum of the intensity values of all the peptides quantified in that sample. Data points not quantified for a particular peptide were given a peptide intensity value equal to the minimum intensity value quantified in the sample divided by 10. Significant differences in phosphopeptide intensities were assessed by student’s t test with Benjamini Hochberg multiple testing correction (false discovery rate).

### Kinase-Substrate Enrichment Analysis (KSEA)

Kinase-substrate enrichment analysis (KSEA) was conducted as previously described by Casado *et al.* (2013). Briefly, phosphopeptides with a p<0.05 (assessed by t test of log2-transformed data) were grouped into substrate sets based on the PhosphoSite database. The ‘enrichment’ method was used to infer differences in the abundance of substrate groups across samples. Z-score was calculated using the formula (Z-score = (mS-mP)*m^1/2^/δ) where m is the size of the substrate group and δ is the standard deviation of the mean abundance across the whole dataset. Z-score was converted to a p value in Excel.

The mass spectrometry phosphoproteomics and proteomics data generated during this study have been deposited to the ProteomeXchange Consortium via the PRIDE partner repository with the dataset identifier PXD023674.

### Immunohistochemistry and scoring

Formalin fixed paraffin embedded (FFPE) mouse tumours were sliced into 4um-thick sections. Slides were baked overnight at 57C prior to dewaxing. Afterwards slides were put into xylene, 100% ethanol and in 3% H2O2/Methanol solution for 10 minutes for endogenous peroxidase inactivation. Antigen retrieval for Ki67 and Cleaved caspase-3 was done using a 10mM citrate buffer (pH 6) for 10 minutes at high power (700W) in a microwave. Samples were left at RT for 30 mins and blocked in 10% goat serum for 20 minutes. Primary antibodies (Ki67 Abcam ab16667, Cleaved caspase-3 Cell signalling #9661) were applied at a 1:100 dilution and left incubating 1h at RT (Ki67) or overnight at 4C (Cleaved caspase-3). Anti-rabbit biotinylated secondary antibody (Vector laboratories BA-1000-1.5) was added at a 1:200 dilution for 30 minutes at RT. Samples were incubated with ABC reagent (Vector laboratories PK-6100) for 20 minutes at RT. Afterwards slides were developed with DAB solution (Dako K3468) for 5 minutes. Washes with PBS (2x, 2 minutes each) were carried out every time slides were incubated with a different reagent.

Finally, samples were counterstained with Haematoxylin, differentiated in 1% acid alcohol, dehydrated in 70%, 90% and 100% ethanol, cleared in xylene and mount using DPX mounting medium. Slides were imaged using the PANNORAMIC 250 Flash III scanner (3D Histech).

For the scoring, ten 50x fields were selected at random so that the tumour regions within the xenograph were uniformly covered. Cleaved caspase-3 stained samples presented a highly apoptotic area at the interphase of the tumour and the stromal regions. Cleaved caspase-3 staining was strong in this area for all samples (regardless of the cell type or the treatment) and was therefore, excluded from the quantifications. For Ki67, the quantification fields were imported into QuPath software and the positive cell detection feature was used to detect the total number of cells, the number of DAB positive and negative cells. For Cleaved caspase-3, ImageJ was used and positive Cleaved caspase-3 cells were manually counted using the Cell Counter plugin. Early and late apoptotic cells were counted. Criteria for considering a Cleaved caspase-3 positive cell was that the DAB signal should show a defined shape and colocalise with Hematoxylin (presence of a nucleus). In some cases, a cell was included where more than one nuclei was observed (late apoptosis).

### Subcutaneous Xenograft study

6 weeks old female NOD-SCID mice were purchased from Charles River and housed with food and water ad libitum, five animals were kept in each cage. A549 LIMD1^−/−^ and LIMD1^+/+^ were grown until reaching the beginning of the exponential growth phase and then detached and resuspended in Matrigel 5mg/ml (Sigma). 1×10^6^ cells in 100 μl of Matrigel were injected subcutaneously into the mice, 20 with A549 LIMD1^−/−^ and 20 with A549 LIMD^+/+^. Tumour growth was measured every two days using callipers and the tumour size calculated using the formula V = (length^2^ x width)/2. Once the average tumour size was between 150 and 200mm^3^, each group was further divided into two groups and once a week received either PF-477736 (7.5mg/kg per dose) or vehicle twice in a day with 6 hours difference. The vehicle contained 50 nM sodium acetate and 4% dextrose, pH 4 (Sigma). Tumour size and mice weight were monitored three times per week and the experiment was stopped when the tumour size exceeded 1.44cm^3^. Mice were then culled and the tumours harvested, sectioned longitudinally and each section was fixed in 10% neutral buffered formalin.

### Statistical analysis

Data was normalized to relevant controls as required. Statistical analysis was conducted using GraphPad Prism 8.0, using the appropriate statistical test for the number of groups and type of data generated from the experiment. Figure legends detail statistical tests used in individual experiments. Statistical significance is shown using the following nomenclature: ns p>0.05, *p≤0.05, **p≤0.01, ***p≤0.001.

## AUTHOR CONTRIBUTIONS

KMD, PG, MFCG, KSB, MH, KS, FKM, MS, RB, PCI, PRC, SAM and TVS designed and performed experiments and analysed the data. All authors contributed to editing and proofreading of the manuscript. KMD, PG, SAM and TVS wrote the manuscript. SAM and TVS supervised and managed all research.

## ACKNOWLEDGMENTS

We thank Julian Blagg, Michelle Garrett, Paul A. Clarke and Dimitris Logos for advice on the study.

---

**Figure S1.**
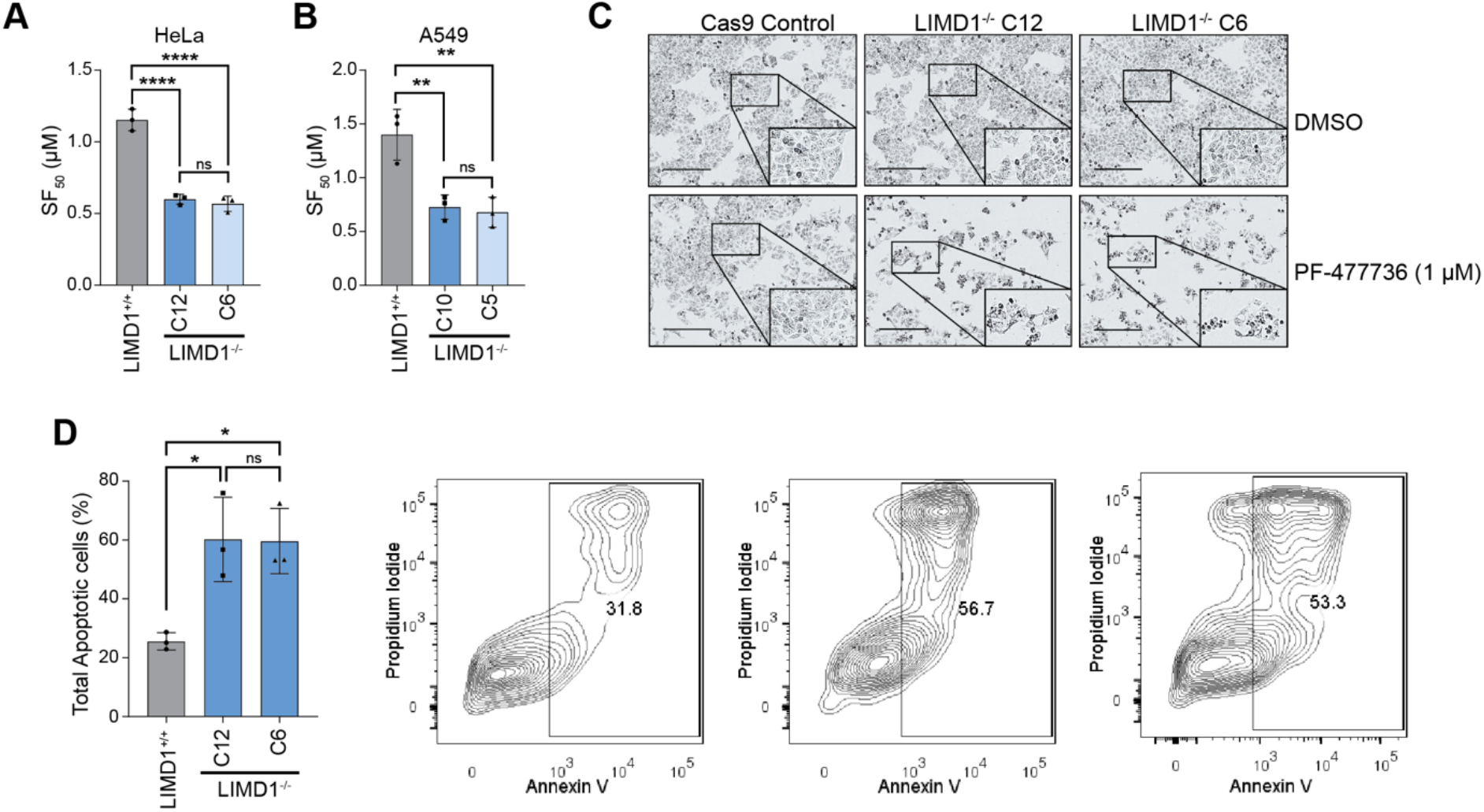
PF-477736 is a selective inhibitor of LIMD1 deficient cells. **A-B)** SF50 values of PF477736 in A549 and HeLa isogenic LIMD1^−/−^ lines (n = 3, one-way ANOVA). **C)** Bright field images of cells upon 48 hours of PF477736 treatment (scale bar = 100 mM, representative images from n = 3). **D)** Bar chart and contour plots of Annexin V/PI staining in HeLa isogenic LIMD1^−/−^ lines. Cells were treated for 48 hours at 1 mM dose of PF477736 before staining and analysis by flow cytometry (n = 3). ns p>0.05, *p≤0.05, **p≤0.01, ***p≤0.001, **** p<0.0001.

**Figure S2.**
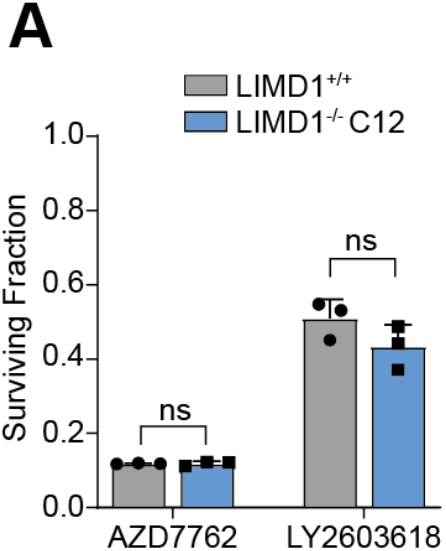
P—477736 selectively kills LIMD1^−/−^ cells independent of Chk1 inhibition. **A)** Alternative Chk1 inhibitors from drug screen in Figure 1A. Surviving fraction determined following 5 days treatment at 1μM.

**Figure S3.**
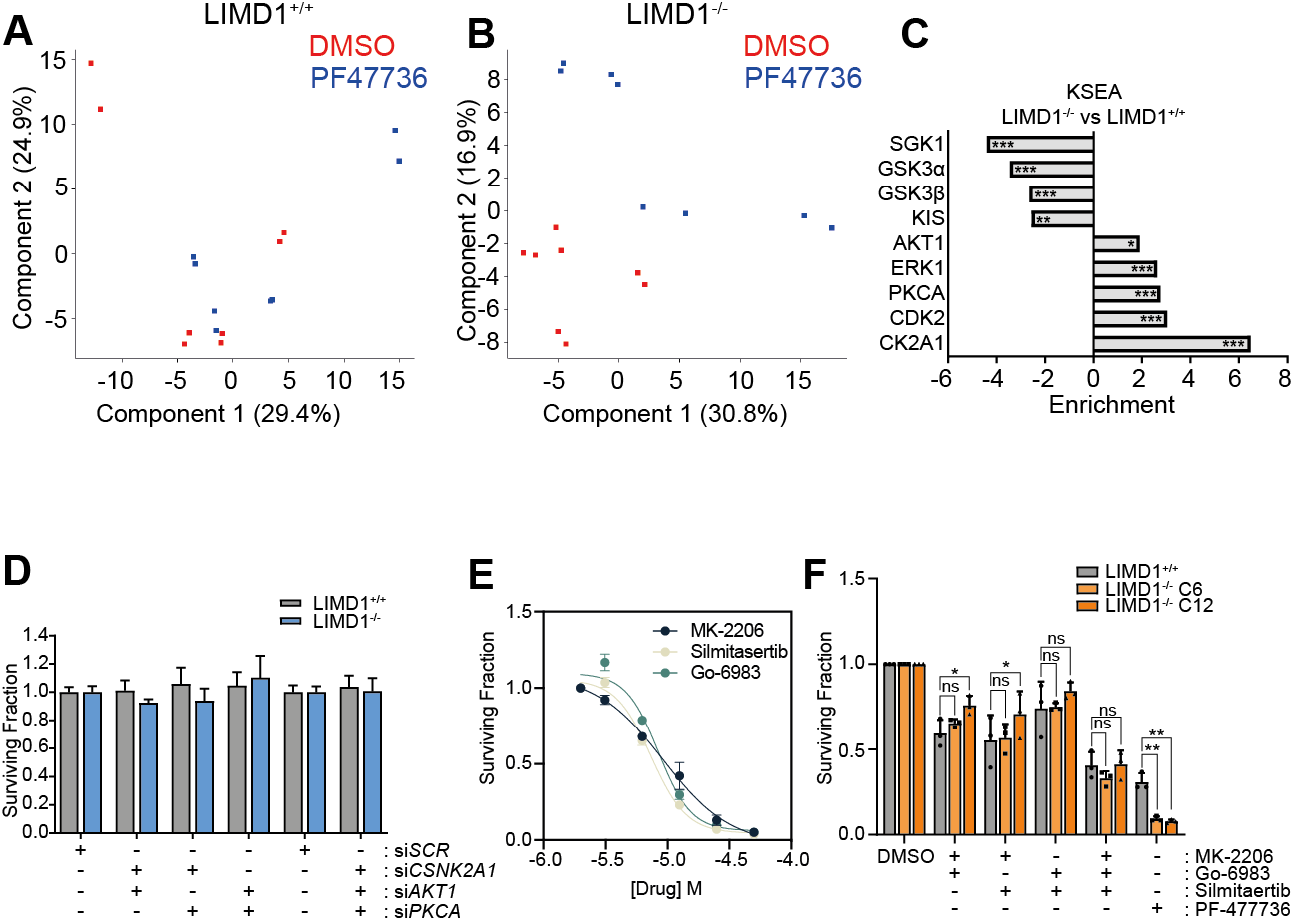
PF-477736 is a broad-spectrum kinase inhibitor that elicits LIMD1^−/−^ specific cellular changes in the phosphoproteome. **A-B)** PCA analysis on phosphoproteome changes shows no separation between treated and untreated samples in LIMD1+/+ cells, but a clear separation in LIMD1−/− cells. Cell viability of HeLa isogenic LIMD1^−/−^ lines treated with siRNA against *CSNK2A1, AKT1* and *PKCA.* Viability was measured 120 hours post transfection and surviving fractions were calculated (n = 3). **B)** Dose response curves of indicated inhibitors in HeLa LIMD1^+/+^ line (n = 2). **C)** Cell viability of HeLa isogenic LIMD1^−/−^ lines treated with inhibitors in combination at their SF_80_ values based upon dose response curves in panel B and PF-477736 at 1 mM as positive control (n = 3, two-way ANOVA). ns p>0.05 *p≤0.05, **p≤0.01, ***p≤0.001

**Figure S4.**
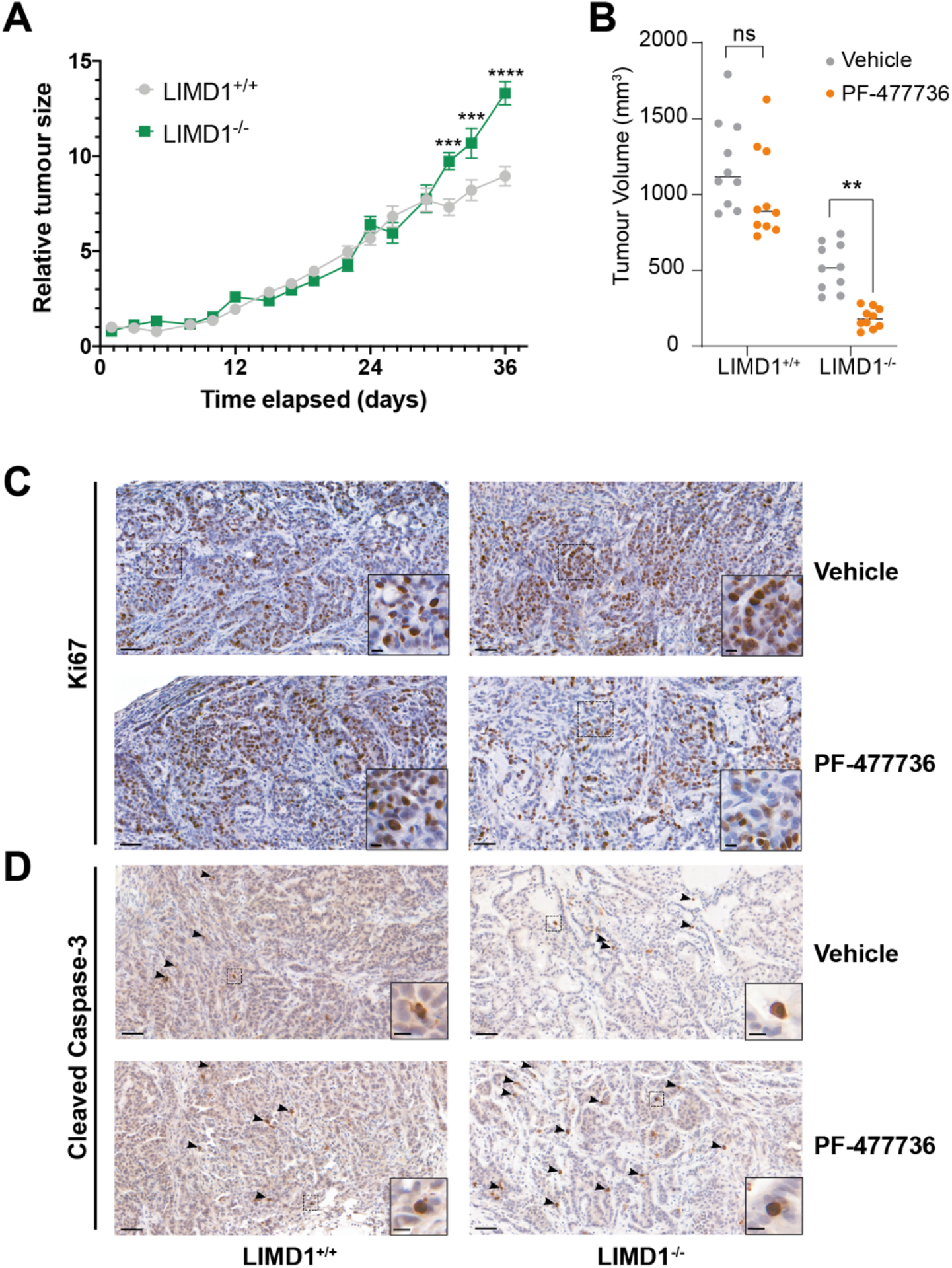
PF-477736 treatment is proof-of-concept inhibitor of LIMD1 deficient lung cancers. **A)** Relative tumour size of vehicle treated xenografts from Fig. 4E normalized to first tumour measurement (n = 10 per group, two-way ANOVA). **B)** Tumour volume of xenografts at day 29 (n = 10 per group, two-way ANOVA). **C-D)** Example staining of Ki67 and Cleaved Caspase-3 in xenografts. Arrows in panel D indicate cleaved caspase-3 positive cells.

